# Inactivation of Airborne Pathogen Surrogates by Triethylene Glycol

**DOI:** 10.1101/2025.08.08.669335

**Authors:** Grishma Desai, Emanuel Goldman, William Jordan, Jamie Balarashti, Jack Caravanos, Rachel Edgar, Etienne Grignard, Gurumurthy Ramachandran, Gediminas Mainelis

## Abstract

The COVID-19 outbreak brought to the fore the importance of airborne transmission in spreading human infectious diseases and highlighted the need for sustainable mitigation strategies. Triethylene glycol (TEG) has been documented as having microbicidal capabilities and has been proposed as one such mitigation strategy. Aerosolized TEG exhibits antimicrobial activity against airborne microorganisms; Grignard Pure™ Technology was developed to safely aerosolize TEG for decontamination of enclosed spaces. Here we show that this TEG formulation effectively inactivates airborne microorganisms, resulting in 2 to 4.5 net log reduction in concentration of viable bacteria, viruses, and mycobacteria within 30-60 minutes at TEG concentration (aerosol + vapor) of ∼0.7 mg/m^3^, which is well within the range considered safe for humans. Our data also demonstrate that aerosolizing both the test organisms and the antimicrobial product provides a more accurate and relevant measure of the product’s efficacy for indoor usage than traditional surface - or solution - based disinfection assays. Accurate evaluation of antimicrobial efficacy is a crucial step in adopting novel interventions and tools to control airborne pathogens that pose a public health risk. Our findings argue that testing protocols must match the intended use of any intervention. Given the safety concerns of aerosolizing human pathogens for direct testing of airborne infectious burden, we also advance an approach for selecting suitable surrogate microorganisms based on their phenotypic and biophysical similarity to corresponding pathogenic species.

**Importance:** During the COVID-19 pandemic, personal protective equipment, social distancing, and even vaccinations proved sub-optimal in controlling the spread of COVID-19. Public health practice and the hierarchy of controls emphasize primary prevention, whereby the pathogen is removed or destroyed before exposure to the public. Triethylene glycol (TEG) has the potential to inactivate airborne pathogens and limit their spread. TEG is designated a “safer chemical” by the US EPA and has been used for decades in aerosol deodorizers and theatrical special effects. This study shows that aerosolized TEG is highly effective at eliminating a wide spectrum of viable airborne pathogen surrogates at concentrations well below the threshold of safety concern. Thus, it may afford significant protection against the transmission of infectious agents with pandemic potential.

## INTRODUCTION

Airborne disease transmission significantly affects global health, economies, and societal structures. Due to their global impact, COVID-19, influenza, and tuberculosis (TB) are probably the best-known examples of airborne diseases. To date, COVID-19 has led to over 7 million fatalities globally (https://data.who.int/dashboards/covid19/cases). Seasonal flu affects millions of people each year, with the World Health Organization (WHO) estimating between 290,000 and 650,000 annual deaths (https://www.who.int/news-room/fact-sheets/detail/influenza-(seasonal)). Additionally, TB continues to be among the leading causes of death worldwide, with around 1.25 million deaths reported in 2023 according to WHO data (https://www.who.int/news-room/fact-sheets/detail/tuberculosis).

The COVID-19 pandemic prompted a re-evaluation of the importance of airborne transmission as a driver of infections, and therefore the interventions needed to combat them (1). When an infected person coughs, sneezes, speaks, or breathes, respiratory pathogens are exhaled within droplets on a size continuum ranging from sub-micrometer to millimeters in size (2). There is an emerging consensus that inhalation of these infectious particles can transmit disease over both short and long distances, and that many factors beyond droplet size determine airborne dissemination (3). The airborne route is likely the dominant mode of transmission for many respiratory diseases, rather than direct deposition of large droplets onto the eyes, nose, or mouth of a susceptible individual, and may be accelerated in enclosed spaces where infectious particles tend to be concentrated (1). In addition to exhalation, infectious agents can disperse into the air from other sources like toilet flushing (e.g. gastrointestinal pathogens such as norovirus), skin blisters (e.g. measles, chickenpox, *Staphylococcus aureus*), or release of fungal spores (e.g. *Aspergillus*) (4)

Public health strategies to curb the spread of airborne diseases typically include practices like the use of face masks and other personal protective equipment, social (physical) distancing, and hand hygiene. Although these strategies can help, they rely on individual compliance. Vaccination is another pillar of infection control, but development or adoption of existing vaccines requires time, and the primary goal is to protect the population from severe disease rather than preventing transmission; people vaccinated against COVID-19 who became infected could still efficiently transmit SARS-CoV-2, for example (5). Ultimately, reducing the infectious burden within the environment will reduce infection risk, whether through improved ventilation, pathogen-neutralizing technologies, or adjusting indoor climate (6). An ideal transmission-blocking intervention should be safe, cost-effective, scalable, environmentally responsible, pan-microbial with minimal chance of resistance development, and compatible with other infection control measures for a layered approach. To have the greatest impact, an intervention should be suitable for long-term proactive use and rapid global deployment early in a pandemic. Low molecular weight non-toxic glycols meet these criteria, and the effectiveness of triethylene glycol (TEG) against various airborne microbes is the focus of this paper.

Microbicidal properties of TEG were identified over 70 years ago, and TEG was identified as a substance with activity against airborne biological threats (7). The microbicidal effects of TEG occur when a sufficient quantity of TEG vapor condenses on particles or droplets carrying microbes (8), which leads to their quick inactivation (the specific mechanism of action is discussed below in this paper). A concentration as low as 2 – 5 mg/m^3^ of TEG in the air was found to be enough for achieving "maximum germicidal action" against a range of airborne infectious agents (7). These concentrations are well below the threshold of safety concern (9). Extensive research in the 1940s and 1950s showed that TEG vapor effectively neutralizes viruses or bacteria responsible for influenza, meningopneumonitis, and psittacosis (10) as well as pneumococci types I, II, and III, beta hemolytic streptococci groups A and C, staphylococci, influenza bacilli, *Escherichia coli*, and *Bacillus aerogenes* (11), and other microbes. However, interest in such primary prevention methods declined as antibiotics became more widely available and effective.

TEG received renewed interest during the COVID-19 pandemic as a potential tool to rapidly inactivate the airborne SARS-CoV-2 virus, thus limiting its spread. In a recent study, we have shown that TEG achieves a net 3-log inactivation within 15 minutes against airborne bacteriophage MS2, a surrogate for viral pathogens, including SARS-CoV-2 (12). Since MS2 is a non-enveloped virus, we speculate that TEG would be even more efficacious against enveloped viruses, including SARS-CoV-2. A related compound, propylene glycol (PG), has also been shown to inactivate several airborne respiratory viruses (13). However, PG requires higher concentrations than TEG, as was observed in the 1940s (8). The antimicrobial activity of various glycols has recently been reviewed (14). The efficacy of TEG against airborne MS2 at different building operational parameters was also recently investigated (15). Thus, evidence indicates that TEG vapor can be used to inactivate airborne microorganisms, thus potentially reducing disease transmission *via* airborne route.

While recent inactivation studies focused on MS2, the application of vaporized TEG as a public health tool for infection control requires this compound to be able to inactivate a variety of airborne microorganisms. This study fills this data gap by demonstrating the broad-spectrum efficacy of TEG against Gram-negative and Gram-positive bacteria, enveloped and non-enveloped viruses, a mycobacterium, and a mold species. The study developed protocols to accurately assess the potential of TEG vapor to inactivate microorganisms in the air, thus enabling a comparison of the microbicidal activity during the intended airborne phase application with conventional surface disinfection testing protocols. The results of our study demonstrate and highlight the need for testing protocols that reflect the ‘real world’ application of novel infection control measures, and the need for appropriate pathogen surrogates that can be safely used to comprehensively assess the control of airborne transmission. Otherwise, surface-centered testing (as is typically performed) obscures the ability of an airborne phase product like TEG to inactivate microorganisms that are primarily infectious via the airborne route.

## MATERIALS AND METHODS

### Test microorganisms

The selection of test microbes was guided by biosafety constraints associated with aerosolized pathogen work. Many clinically relevant organisms, such as *Staphylococcus aureus*, require BSL-2 containment for benchtop procedures and BSL-3 or higher for aerosolized studies. As most commercial and academic aerosol testing laboratories, including the lab utilized for these studies, are equipped with only BSL-2 aerosol chambers, testing in the airborne phase is limited to BSL-1 organisms. Biosafety guidelines, such as the CDC BMBL (6th Edition), explicitly note that additional primary containment and personnel precautions—such as those recommended for BSL-3—should be considered for activities with a high potential for aerosol or droplet production (16). Similarly, the WHO Laboratory Biosafety Manual (3rd Edition) emphasizes that aerosol-generating procedures may require a higher biosafety level than that indicated by an organism’s Risk Group, based on risk assessment (17). Therefore, appropriate BSL-1 surrogates, based on the similarity of the phenotypic traits, were selected to assess the response of corresponding BSL-2 pathogens to treatment by TEG.

This study prioritized diverse microbial species, representing a broad spectrum of potential pathogens. The study aimed to evaluate the antimicrobial agent’s broad-spectrum potential under realistic conditions by incorporating organisms with varying susceptibilities and structural differences. The organisms selected for the study included:

- Gram-negative bacteria: *Pseudomonas fluorescens* (ATCC 13525) (American Type Culture Collection, Manassas, VA), *Salmonella typhimurium* (ATCC 53648), and *Klebsiella aerogenes* (ATCC 51697).
- Gram-positive bacteria: Methicillin-resistant *Staphylococcus epidermidis* (ATCC 12228) and *Listeria innocua* (ATCC 33090).
- Viral species: Non-enveloped bacteriophage MS2 (ATCC 15597-B1) and enveloped bacteriophage Phi6 (ATCC 21781-B1).
- Acid-fast Gram-positive bacterium: *Mycobacterium smegmatis* (ATCC 607).
- Mold species: *Aspergillus brasiliensis* (ATCC 16404).

The microbes above were exposed to airborne TEG in the airborne phase, and to liquid TEG while surface-bound to carriers.

In another subset of experiments, we evaluated the inactivation of surface-bound actual pathogenic strains and their Biosafety Level 1 (BSL-1) surrogate counterparts (i.e., *S. epidermidis, K. aerogenes*, *P. fluorescens*) in parallel when exposed to TEG vapor. This test also advanced the selection of appropriate surrogates, providing justification for their use in evaluating airborne antimicrobial efficacy of TEG.

The pathogenic species were:

- Gram-positive bacteria: methicillin-resistant *Staphylococcus aureus* (ATCC 6538)
- Gram-negative bacteria: *Klebsiella pneumoniae* (ATCC 4352), Gram-negative bacteria: *Pseudomonas aeruginosa* (ATCC 15442)

### Inoculum preparation

Pure strain seed stocks were sourced from the ATCC. Working stock cultures were prepared under aseptic conditions using a Class II biological safety cabinet (Labconco Purifier Class II Safety Cabinet, Kansas City, MO), following ATCC guidelines. Strains were revived in appropriate growth media and incubated under conditions specific to each organism, following ATCC guidelines. Except for *Listeria innocua*, all other bacterial species were cultured in tryptic soy broth and plated on tryptic soy agar. *Listeria innocua* was cultured in brain heart infusion broth and plated on brain heart agar. *Aspergillus brasiliensis* was used as a dry spore powder and plated on potato dextrose agar. Serial dilutions were performed in sterile phosphate-buffered saline (PBS) supplemented with 0.005% Tween 80 to facilitate homogeneity.

For virus strains, the host bacteria (*E. coli* for MS2 and *P. syringae* for Phi6) were grown overnight in tryptic soy broth at 37°C for *E. coli* and 28°C for *P. syringae*. During the logarithmic growth phase, the bacterial suspension was infected with either MS2 or Phi6 bacteriophage. After 24 hours, the infected cells were lysed, and cellular debris was removed by centrifugation at 3000 rpm for 15 minutes. The resulting supernatant was poured off from the cellular debris, sampled, serially diluted, and plated to determine the viral concentration.

The stock concentrations of test microorganisms were verified through colony-forming unit (CFU) or plaque forming unit (PFU) enumeration. Stock dilutions with microorganism concentrations ranging from 10^9^ to 10^10^ CFU/ml or PFU/ml were used for aerosolization, as described below.

### Sample processing

In the inactivation experiments described below, the airborne test microorganisms were directly captured into PBS using impingers, while the surface-bound organisms on glass coupons were eluted into PBS using 50ml conical tubes (Celltreat, Pepperell, MA), followed by enumeration by culturing in triplicate. If necessary, the collected samples were diluted by a series of ten-fold dilutions before culturing. Appropriate dilutions of virus samples underwent small-drop plaque assays technique, while bacterial and mold spore samples were plated using a spread plate technique. *Listeria innocua* was cultured on Tryptic Soy Agar supplemented with 5% sheep blood, *Aspergillus brasiliensis* was grown on Potato Dextrose Agar with chloramphenicol, and *Mycobacterium smegmatis* was cultured on Middlebrook 7H11 Agar. Tryptic Soy Agar was used for all other species. All media were supplied by Hardy Diagnostics, Santa Maria, CA. All samples were plated across a minimum of three dilution ranges, with triplicate plating per dilution range on pre-labeled, culture plates with species-appropriate agar and organism. Plates were incubated at species-specific temperatures ranging from 28–37°C for 24–48 hours before enumeration. Counts between 5–50 colony-forming units (CFU) or plaque-forming units (PFU) per plate were recorded; counts outside of this range were not counted. This approach ensured consistent quantification across all tested conditions.

### Inactivation of airborne microbes by airborne TEG (aerosol and vapor)

Airborne inactivation studies were conducted using paired trials for each microbial species discussed in this paper. For every species, a positive control trial and a corresponding inactivation trial were performed using a single, unified biological stock that was split between the two conditions. This approach ensured both trials began with identical starting material. All other trial conditions—such as the nebulizer type, nebulization pressure and duration, hold and sampling times, and methods for collection, elution, and plating—were kept constant between the paired trials to ensure consistency.

During the study, airborne concentrations of viable test microbes were measured at different time points for each paired control with and without exposure to the inactivating agent, triethylene glycol (TEG). TEG was aerosolized through Grignard Pure™ Technology (Bleu Garde LLC, Rahway, NJ; https://www.bleugarde.com) developed a few years ago to disperse TEG in aerosol and vapor form.

Inactivation of airborne microbes was investigated in an insulated ∼16 m³ stainless steel (SS 304) bioaerosol chamber with internal dimensions 9.1 ft × 9.1 ft × 7 ft (2.8 x 2.8 x 2.1 m), with a total volume of just over 16,000 liters. It features three windows and a hermetically sealed door to maintain environmental integrity. Environmental parameters, including temperature and relative humidity, were continuously monitored using embedded sensors (HT.w Sensors, SensorPush, Brooklyn, NY) within the chamber. During experiments, the average temperature and RH were 22-24°C and 30-40%, respectively. Aerosol samples were collected through two stainless steel probes (3/8-inch inner diameter) located 18 inches from the chamber’s sidewalls at a height of 36 inches. Sampling points were positioned at the diagonal corners of the chamber to ensure representative samples. Two interior mixing fans, operating at ∼50 cubic feet per minute (cfm) each (equivalent to about 10 internal ACH), were used for the entirety of all trials to provide bulk mixing and to ensure homogeneity of the chamber during the test period. This setup ensured uniform sampling across the chamber’s interior for precise and reproducible results (18).

The timeline of inactivation experiments is shown in Figure 1. For most of the organisms tested, Grignard Pure™ solution in its undiluted form (Lot#021722) was aerosolized via the Aura device (Bleu Garde, LLC), which was positioned in the center of the chamber and activated remotely. *A. brasiliensis* was treated with the Clearify device (Bleu Garde, LLC) due to anticipated need of higher TEG concentration because of spores. First, the chamber was pretreated with aerosolized Grignard Pure™ for 30 minutes (from t = -40 min to t = -10 min in Fig. 1), which simulated an indoor environment treated with Grignard Pure™. After this 30-min pretreatment period, while Aura (or Clearify) dispersal device continued to operate, bioaerosols were aerosolized into the chamber over a 20-min period using a 1-inch diameter stainless steel conduit. Bacteria and viruses were aerosolized using 24-jet Collison nebulizer (CH Technologies, Westwood, NJ) operated at 40 psi. *A. brasiliensis* spores were disseminated via a dry powder eductor constructed from stainless steel and operated using in-house filtered compressed air. Since microorganisms interacted with TEG while being introduced, we designated the mid-point of this 20-min aerosolization period as t= 0 min. Once the bioaerosol introduction was completed, this time marked time t = 10 minutes in our treatment timeline, because that is the average duration that the organisms have interacted with airborne TEG (Fig. 1). Bioaerosol samples were started at time points t = 10, 35, 50, and 80 minutes using two AGI-30 impingers (Ace Glass Incorporated, Vineland, NJ), and sampling duration was 10 minutes. Impingers contained 20 mL of sterile PBS supplemented with 0.005% Tween 80 to enhance capture efficiency and promote microorganism dispersion within collection liquid. The impingers operated at a flow rate of 12.5 L/min, provided by 1/3 HP Gast Rotary Vane Vacuum Pumps (Eureka, MO). After each sampling period, the contents of the two impingers were combined to provide a chamber average and transferred to sterile conical tubes (Celltreat, Pepperell, MA) for microbial enumeration. A single low velocity mixing fan (12” Oscillating Black HTF1220B, Honeywell, Charlotte, NC) was used in the chamber during testing to ensure a homogenous microbe distribution throughout the chamber and promote GP and organism mixing. Between samples, impingers were thoroughly cleaned and sterilized by immersion in a 3% bleach solution, followed by rinses with tap water and distilled water before air drying. Following the experiments, the remaining particles and vapor were evacuated from the chamber (19).

**Figure 1.**
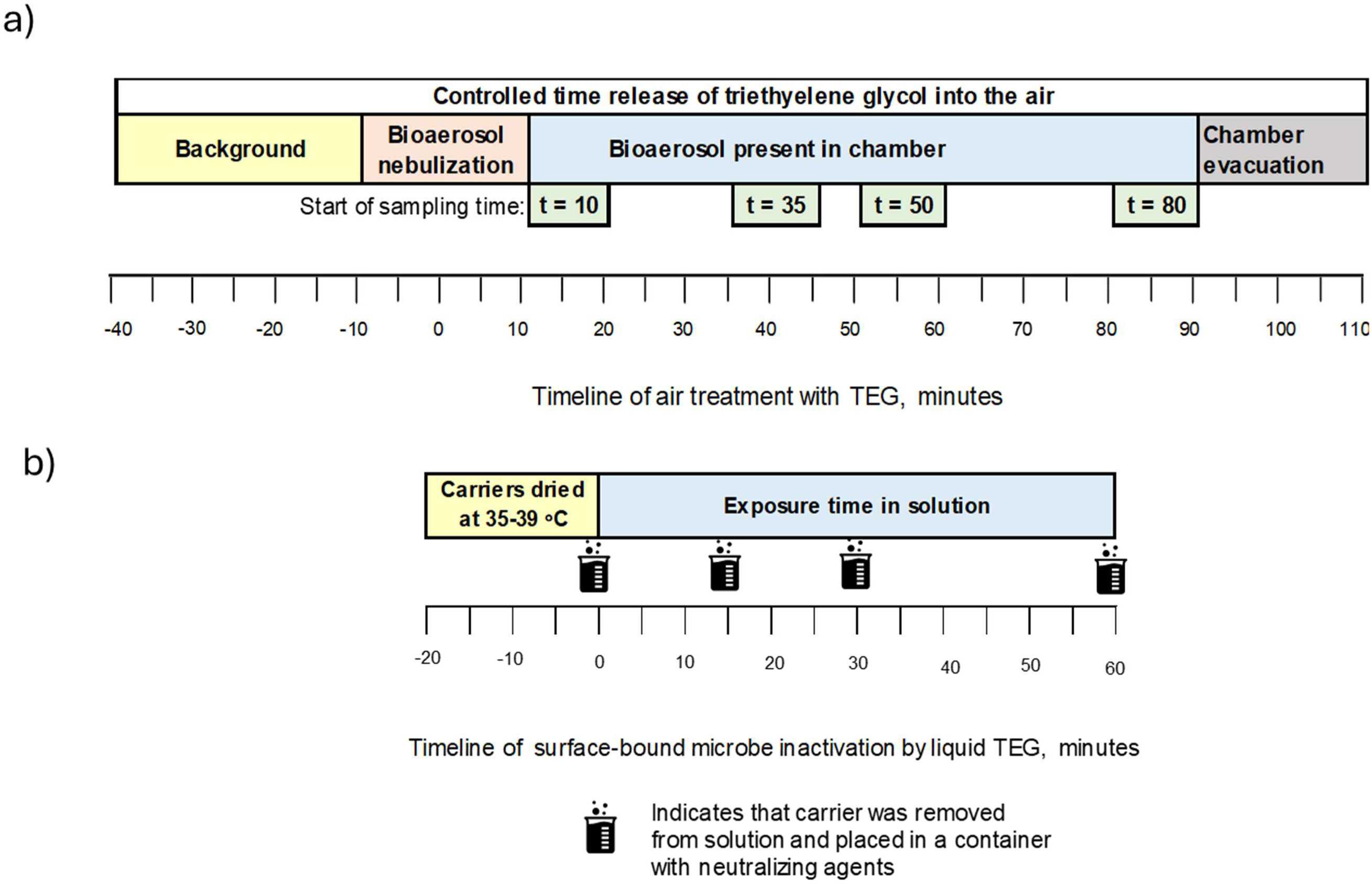
Timeline of experiments investigating the efficacy of triethylene glycol (TEG) against microbes: a) inactivation of airborne microbes and b) inactivation of surface-bound microbes. See text for full description.

The aerosol concentration of Grignard Pure™ was estimated using a calibrated TSI SidePak AM520 (TSI, Inc., Shoreview, MN). The Grignard Pure™ aerosol concentration can be easily converted to TEG aerosol concentration as described in our earlier study (12). The total TEG concentration (aerosol + vapor) was estimated using an earlier-developed correlation between AM520 particle concentration data and NIOSH tube sampling method 5523, which determines the total (aerosol + vapor) airborne concentration of TEG. During the inactivation of most airborne microbes (except *A. brasiliensis*), Aura device was used, and the average TEG aerosol and total concentrations were 0.04 ± 0.02 and 0.66 ± 0.35 mg/m^3^, respectively. In anticipation of higher resistance of fungal spores, for testing with *A. brasiliensis*, we used the Clearify device, and the total TEG concentration was increased to 1.96 mg/m^3^, of which 0.3 mg/m^3^ was in aerosol form. Data from this experiment are provided in Table S1 (Supplemental Material).

In control trials for each test microbe, the procedure was exactly the same as described above, except Grignard Pure™ was not introduced into the chamber. The control trial accounted for the loss of viable microorganisms due to gravitational settling and natural die-off. The efficacy of TEG was determined by comparing the airborne concentrations of viable microorganisms - measured as colony-forming units per m^3^ (CFU/m^3^) for bacteria and plaque-forming units per m^3^ (PFU/m^3^) for viruses - collected at the same time points as during air treatment with TEG. The inactivation efficiency was determined for each sampling point as a percent value and log reduction relative to microorganism recovery without treatment. The initial microorganism concentrations with and without treatment by TEG were normalized. The applied mathematical expressions are presented in our earlier work (12). For all microbes, except MS2, there was a single positive control trial and a corresponding TEG inactivation trial. For MS2, two sets of experiments were conducted, and average values are presented.

### Neutralization Verification

Neutralization verification was performed using procedures aligned with EPA SOP MB-26-03, Standard Operating Procedure for Neutralization of Microbicidal Activity Using the Quantitative Method for Evaluating Bactericidal and Mycobactericidal Activity of Microbicides on Hard, Non-Porous Surfaces (20).

These tests confirmed that microbial inactivation attributed to TEG occurred exclusively during airborne exposure and not due to residual TEG activity post-sampling. To assess this, test organisms were directly exposed to TEG in liquid at defined concentrations and contact times, then plated and enumerated. Results were compared to unexposed controls to determine any residual biocidal activity post-collection.

TEG exposure concentrations were calculated based on known airborne concentrations and impinger sampling volumes. These values were then multiplied by a safety factor of 10× to establish the challenge concentration for neutralization testing. A neutralizer solution consisting of TEG10× + phosphate-buffered saline (PBS) + 0.005% Tween 80 was used; PBS + 0.005% Tween 80 served as the neutralizer control.

Triplicate 50 mL conical tubes were prepared for both the TEG-neutralizer and control solutions, each containing 20 mL. Cultures were diluted to <1.00 × 10³ CFU or PFU/mL before being added to the tubes. After a 15-minute contact time, samples were serially diluted, plated using previously described methods, incubated, and enumerated.

Comparisons were made to PBS-only controls and water-diluted controls to verify neutralization efficacy. ANOVA analysis confirmed that PBS + 0.005% Tween 80 had no statistically significant impact on microbial recovery for any species tested, validating that reported reductions were due solely to in-chamber inactivation and not post-collection residual activity.

### Inactivation of surface-bound organisms by liquid TEG

Currently, the efficacy of most antimicrobial products is typically evaluated using surface-focused methods - such as ASTM E1153 (21), ASTM 1053 (22) and the AOAC use-dilution method (23) – that measure the reduction of microorganisms deposited on hard, non-porous carriers following exposure to a liquid, spray, or wipe. We posited that products intended for inactivation of airborne microorganisms must be evaluated using test methods specifically designed to assess their efficacy against airborne microorganisms, as surface-based tests do not assess this mode of transmission. The comparison of the two modes of inactivation (e.g., microorganisms in the air vs surface-bound) is crucial to properly determine a product’s suitability to act an airborne antimicrobial. Therefore, inactivation of airborne microorganisms by Grignard Pure™ in aerosol and vapor form described above was compared to inactivation of surface-bound microorganisms by liquid Grignard Pure™. The methodology for surface-bound testing adhered to ASTM E1153 (21) guidelines for bacterial and fungal assessments on non-porous surfaces and ASTM E1053-20 (22) for virucidal evaluations. While Grignard Pure™ is not classified as a disinfectant or sanitizer, these standardized protocols provided a structured framework to assess the efficacy of TEG against microbial targets in both aerosolized and liquid forms.

Microbial inocula were prepared using the same procedure as described above for aerosol phase tests; however, to simulate environmental contamination, a soil load consisting of 0.3% bovine serum albumin, 0.3% mucin, and 0.3% yeast extract was added to the microbial stocks in accordance with ASTM E2197 and OECD Guideline 13 standards (24).

The timeline for surface-bound microbe inactivation experiments is shown in Figure 1b. Quadruplicate sets of 304 stainless steel carrier discs (0.8mm thick, 2cm diameter) were inoculated (covered) with 0.03 mL of prepared microbial suspension, ensuring uniform coverage. Carriers were dried at 35–39°C for approximately 20 minutes or until visibly dry. For exposure to TEG, carriers were submerged in 5 mL of the Grignard Pure™ solution for 0, 15, 30, and 60 min. For 0-min exposure, the carriers were dipped into the Grignard Pure™ solution and immediately pulled out. After exposure, 20 ml of PBS + 0.005% Tween 80 was added to the container with the test solution and carrier to halt antimicrobial activity. The container was then vortexed for 2 minutes, followed by shaking for 30 seconds to recover microbes from the carrier surface. The recovered microbes were then cultured onto species-specific media using a standard culturing or small plaque assay across various dilutions and incubated at species-specific temperatures (28–37°C) for 24–48 hours before enumeration. The number of CFU or PFU recovered from exposed carriers were compared with the number of carriers not exposed to liquid Grignard Pure™ but otherwise handled in the same way. These control carriers were exposed to the air for 0, 15, 30, 60 min. The inactivation efficacy was determined as a percentage and net log reduction relative to the unexposed carriers. Data from this experiment are provided in Table S2 (Supplemental Material).

### Inactivation of surface-bound organisms by TEG vapor

As briefly discussed above, the selection of appropriate surrogate microorganisms is important because many laboratories are limited to working with airborne BSL-1 organisms only, yet many regulatory procedures require showing evidence of antimicrobial efficiency against BSL-2 organisms. Therefore, we compared the inactivation of several BSL-1 microorganisms with their BSL-2 counterparts when exposed to TEG aerosol and vapor while surface-bound. Microbial cultures and their carriers were prepared as described above for the testing of inactivation by liquid TEG. The experiments were carried out in the same bioaerosol chamber at an average temperature of 22-24°C and 30-40% RH. The chamber was then pretreated with aerosolized Grignard Pure™ for 30 minutes using the Aura device. TEG concentration in this experiment was 0.54 mg/m^3^. Dried carriers in triplicate were placed on a stainless-steel workbench within the bioaerosol chamber at a distance of approximately 3 feet from the dispersal device. The chamber was sealed, and the Aura dispersal device was operated according to its specifications for a total exposure duration of 90 minutes. After exposure, carriers were transferred to labeled conical tubes containing neutralizing buffer and vortexed for 2 minutes, followed by shaking for 30 seconds to recover microorganisms.

The recovered microorganisms were serially diluted in microcentrifuge tubes containing sterile PBS as diluent and enumerated as described above. The number of CFUs recovered from exposed carriers were compared with the number of CFU from carriers not exposed to TEG in the chamber but handled in the same way. The inactivation efficacy was determined as a percentage and net log reduction relative to unexposed carriers. Data from this experiment are provided in Table S3 (Supplemental Material).

## RESULTS

### Overall inactivation of airborne microorganisms

Inactivation of viable test microorganisms in the air by airborne TEG and on hard surface by liquid TEG as supplied by Grignard Pure™ is presented in Figures 2 and 3. Figure 2 shows data for tested Gram-positive and Gram-negative bacteria, while Figure 3 presents inactivation data for tested bacteriophages, acid-fast microorganism, and a mold. The inactivation data are presented as net log inactivation and also as net percent inactivation for different sampling time points (Figure 1). The raw data are provided in Table S1 (Supplemental Material).

**Figure 2.**
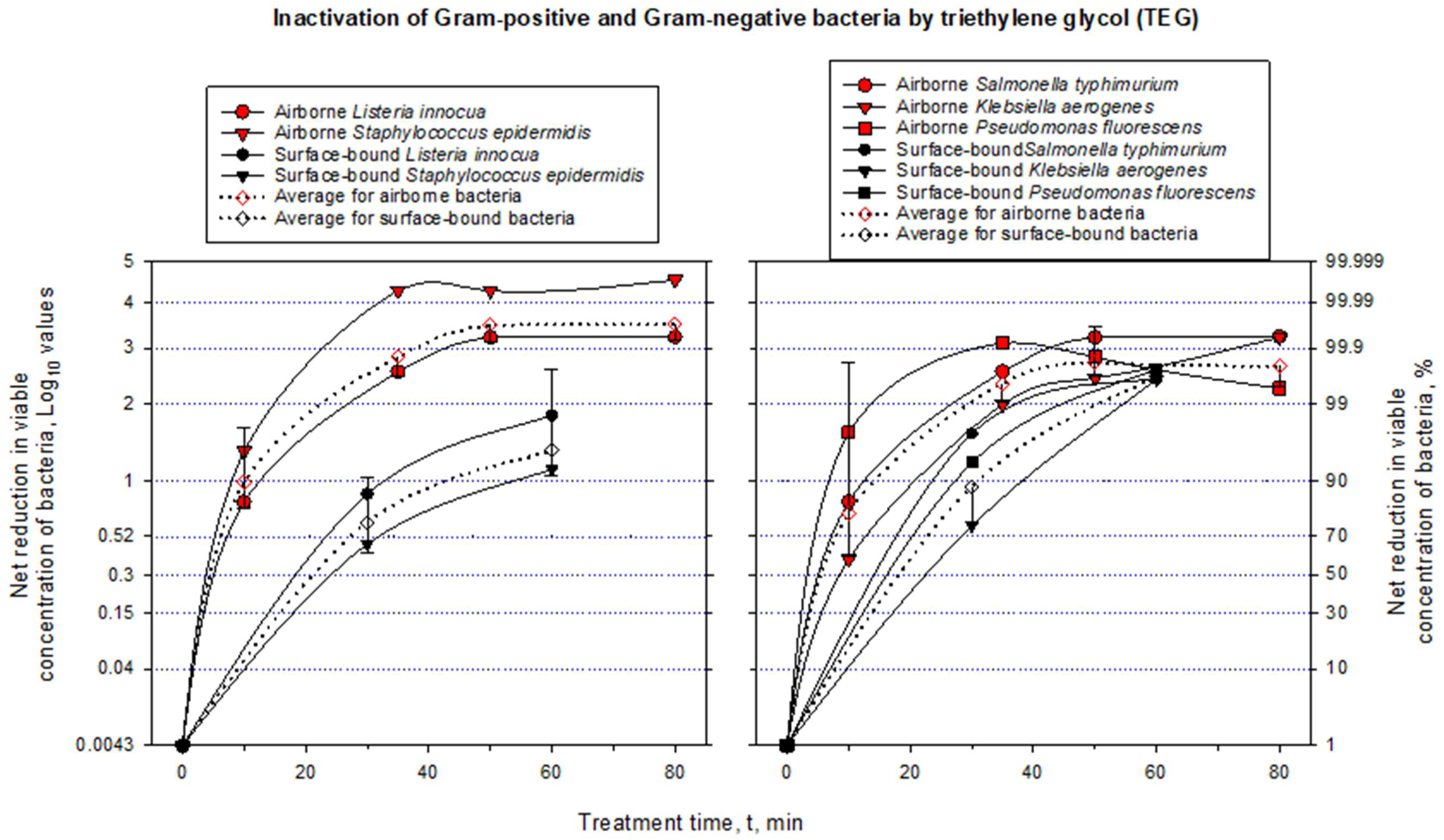
Inactivation of Gram-positive and Gram-negative bacteria by triethylene glycol (TEG). Net log and percent reduction in airborne and surface-bound concentrations of viable bacteria due to treatment by triethylene glycol (TEG). Airborne bacteria were treated by aerosolized TEG, while surface-bound bacteria were treated by liquid TEG. Treatment time for airborne bacteria also denotes the start of 10-min sampling by impingers. See Materials and Methods for experimental details. The propagated error for each data point is provided in Supplemental Materials.

**Figure 3.**
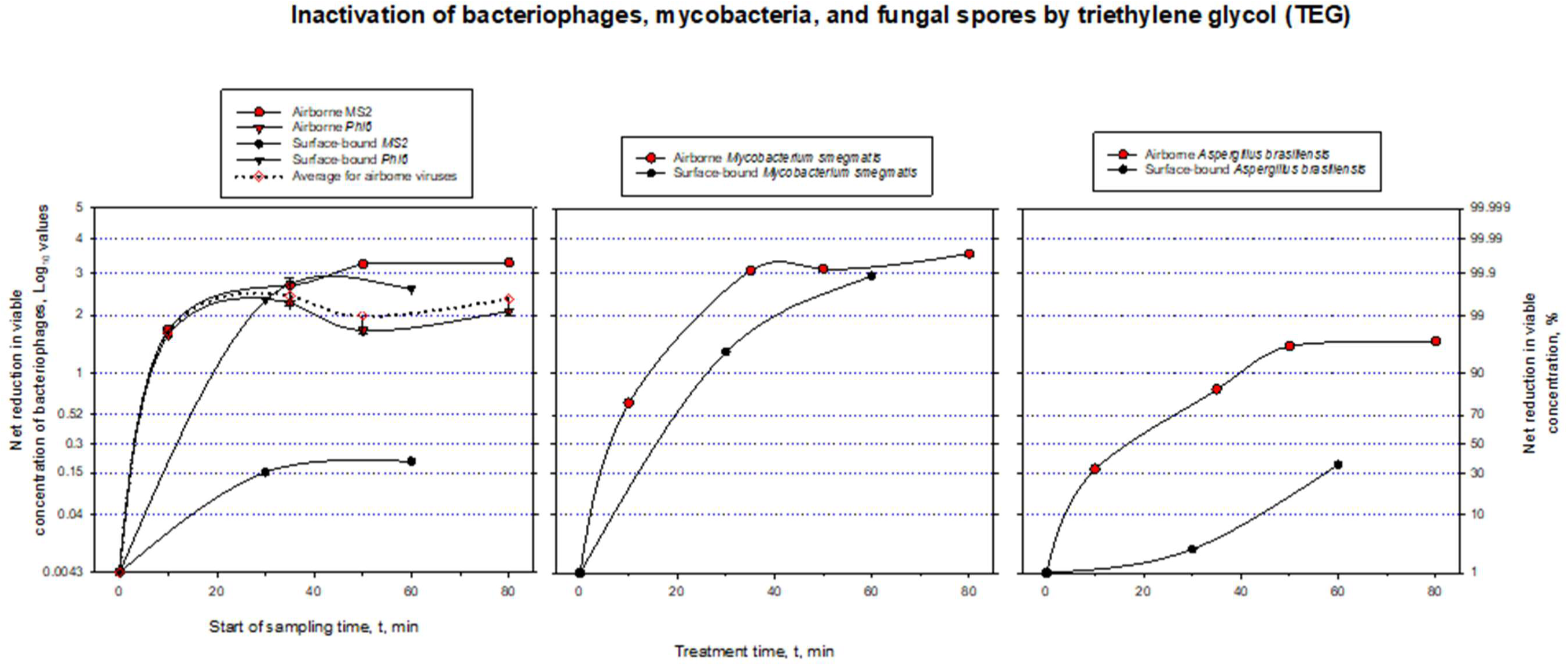
Inactivation of bacteriophages, mycobacteria, and fungal spores by triethylene glycol (TEG). Net log and percent reduction in airborne and surface-borne concentrations of viable bacteriophages, rnycobacteria and fungal spores due to treatment by triethylene glycol (TEG). Airborne microbes were treated by aerosolized TEG, while surface-bound microbes were treated by liquid TEG. Treatment time for airborne microbes also denotes the start of 10-min sampling by impingers. The propagated error for each data point is provided in Supplemental Materials.

Substantial inactivation of all tested airborne microorganisms, except mold, was observed at the 10-minute sampling point, with bacterial inactivation efficacy ranging from 58 to 95%. Both bacteriophages were inactivated with 98% efficacy at the same 10-minute sampling time point. Inactivation of *A. brasiliensis* was 32% at the same time point. All microbes except *A. brasiliensis* were inactivated with at least 99% efficacy at the 35-minute sampling mark. Inactivation of *A. brasiliensis* spores at that time point was ∼80%. At later sampling points, the net inactivation efficacy remained steady or slightly decreased (e.g., Phi6 and *P. fluorescens*) due to increasing natural die-off of the organisms, which led to a reduced net efficacy because the initial concentration of airborne microbes turned out not to be high enough. Both Phi6 and *P. fluorescens* are known to be sensitive to environmental stressors (25, 26), and prohibitively large airborne concentrations of them are needed to observe net inactivation over longer periods. By contrast, inactivation of the same organisms on hard surfaces proceeded much more slowly, with lower inactivation than in the airborne phase at the 30-minute mark for all tested microbes, except Phi6 bacteriophage.

### Inactivation of airborne bacteria

Among airborne bacteria, the highest inactivation was achieved for Gram-positive bacteria, where viable *S. epidermidis* was reduced by 4.28 logs at the 35-minute sampling point, and its inactivation was more than 4 logs at 50 and 80 minutes; *L. innocua* showed more than 3 log reduction at 50- and 80-min sampling points (Fig. 2). When the airborne inactivation data of the two Gram-positive bacteria are pooled, we observe average inactivation of 90% inactivation at the 10-min sampling point and 99% inactivation at all later time points (Figure 2). By contrast, hard surface treatment resulted in slower viability reductions, with *L. innocua* and *S. epidermidis* inactivated by less than 1 log by 30-min treatment and less than 2 logs by 60-min treatment. A similar dynamic was observed for all Gram-negative bacteria. Treatment of airborne organisms again proved superior, with *P. fluorescence* reduced by 3.10 logs at 35 minutes, followed by *S. typhimurium* (2.5 logs) and *K. aerogenes* (2 logs). At later sampling points, their inactivation stayed the same or slightly decreased (e.g., *P. fluorescens*) due to the role played by the natural die-off and settling in inactivation calculation. The pooled airborne inactivation data from three Gram-negative organisms show an average inactivation of 80% at the 10-min sampling point and 99% inactivation at later sampling time points (Figure 2). Viability reduction on hard surface by liquid TEG was notably slower but eventually reached about 2.5 logs for all three bacteria by 60 minutes. A comparison of averaged surface-bound bacteria inactivation data with averaged airborne bacteria inactivation data emphasize the more rapid and significant inactivation of airborne bacteria by airborne TEG compared to surface treatments by liquid TEG.

### Inactivation of airborne viruses, mycobacteria, and fungi

Figure 3 shifts focus to the inactivation of bacteriophages. Aerosol treatment of both bacteriophages achieved a rapid viability reduction: ∼98% inactivation at the 10-min sampling mark. At other sampling points, MS2 was inactivated with >99.9% efficacy, and inactivation of Phi6 varied between 98% (50 min) and 99.5% (35 min). The averaged data for the two bacteriophages show 98% inactivation at the 10-minute sampling point and at least 99% for later sampling time points. The surface inactivation of the two bacteriophages proceeded differently. The reduction of MS2 on a hard surface proceeded slowly, reaching only a 0.20-log (37%) in 60 minutes. By contrast, due to the sensitivity of Phi6 to environmental stressors, its inactivation on the surface was also relatively rapid, reaching 2.2 logs (99.5%) at the 30-minute mark and exceeding that in the airborne state at the 60-minute mark.

Airborne *M. smegmatis* was also rapidly inactivated: by 3-logs at the 35 min mark, and then the inactivation reached a plateau of 3.54 log by 80 minutes. Inactivation of this bacterium on the hard surface proceeded substantially slower, but it reached 2.93-logs by 60 min mark. Its inactivation profile in the air was slower compared to Gram-positive bacteria, but similar to that of some Gram-negative bacteria.

Mold spores are known to be resistant to environmental stress, but airborne TEG achieved 1.5-log inactivation of airborne *A. brasiliensis* after 50 min, and the level stayed the same after 80 minutes of interaction. Inactivation of these spores on the surface was much slower and reached only 0.19-log reduction in 60 minutes. *A. brasiliensis* appears to be more resistant overall compared to tested bacteria, for both airborne and surface testing. It is also worth mentioning that the airborne TEG concentration when inactivating *A. brasiliensis* was almost three times higher than when treating other organisms, confirming the anticipated hardiness of this species. Nevertheless, about 95% of the airborne spores were inactivated after 50-min treatment by airborne TEG.

### Summary of airborne inactivation results

Taken together, Figs. 2-3 demonstrate that TEG provided by Grignard Pure™ is highly effective in reducing the presence of a wide range of viable organisms, particularly in airborne environments where aerosol application consistently outperforms hard surface treatments by liquid Grignard Pure™ in both speed and magnitude of reduction. The findings highlight its robust efficacy against bacteria such as *S. epidermidis* and *S. typhimurium* as well as against bacteriophages like MS2 and Phi6, while also confirming its effectiveness against more resilient organisms like *M. smegmatis* and *A. brasiliensis* over time. These results underscore the utility of Grignard Pure™ in antimicrobial strategies aimed at controlling airborne pathogens in enclosed diverse settings, while offering complementary benefits for surface decontamination over longer durations when applied in liquid form.

### Comparison of inactivation of pathogens with their corresponding surrogates

The next set of experiments examined the inactivation of a select set of BSL-1 organisms and their BSL-2 counterparts when first deposited on slides and then exposed to TEG vapor and droplets for 90 min (Fig. 4). The selected BSL-1 organisms serve as surrogates for their BSL-2 counterparts. Among the BSL-1 surrogates, *K. aerogenes* demonstrated the highest reduction (0.49 log), closely mirroring its BSL-2 counterpart *K. pneumoniae* (0.46-log reduction). This similarity confirms the suitability of *K. aerogenes* to be used as a surrogate for pathogenic *K. pneumoniae*. Likewise, inactivation of *P. fluorescens* (0.18 logs) was similar to that of BSL-2 *P. aeruginosa* (0.21 logs), further validating its use as a representative model for air testing studies. Both Gram-positive species, BSL-1 *S. epidermidis* (0.17-log reduction) and BSL-2 *S. aureus* (0.14-log reduction), showed lower reductions compared to other species but maintained consistent responses between the surrogates and pathogenic counterparts. Overall, a paired t-test of all three microorganism pairs showed that there was no statistically significant difference (p>0.05) among inactivation of the BSL-1 bacteria (surrogates) and BSL-2 bacteria (pathogens). This consistent response between the BSL-1 bacteria and their BSL-2 counterparts demonstrates the relevance and suitability of selected surrogates in assessing treatment efficacy against higher-risk BSL-2 pathogens under safe laboratory conditions. Fig. 4 highlights how closely surrogate organisms replicate the behavior of their pathogenic counterparts under similar exposure conditions. Notably, Gram-negative *K. aerogenes* and *P. fluorescens* provide reliable models for airborne pathogen testing due to their similar responses to *K. pneumoniae* and *P. aeruginosa*, while the same can be said about Gram-positive *S. epidermidis* and *S. aureus*.

**Figure 4.**
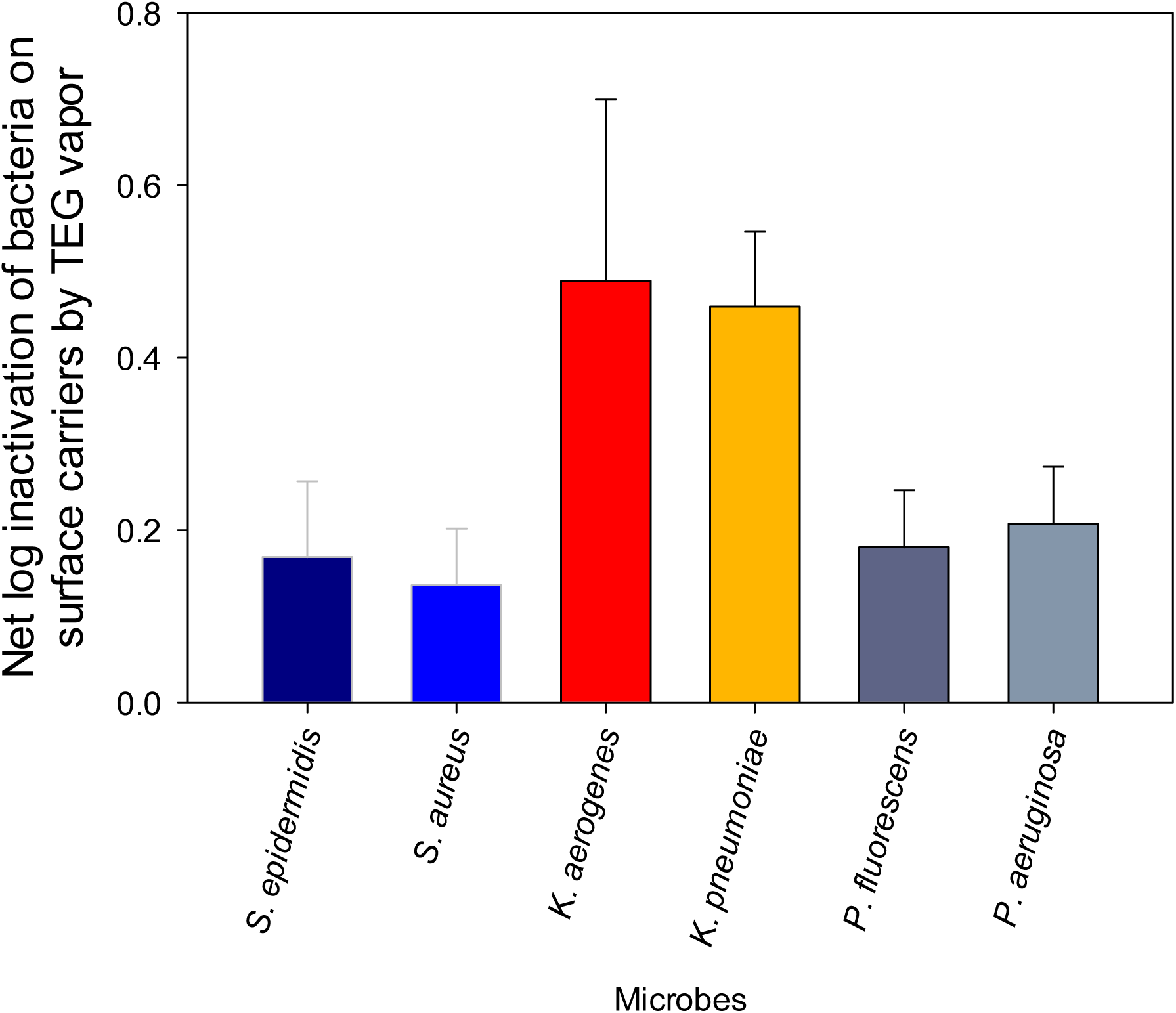
Comparison of inactivation by TEG of bacterial pathogens and their non-pathogenic surrogates. Net log inactivation of surface-bound bacteria by TEG vapor as delivered by Grignard Pure™. The data are mean values of four replicates (carriers) and error bars represent one standard deviation.

## DISCUSSION

### Efficacy Testing Protocol

Although standardized methods exist for evaluating the efficacy of antimicrobial products in liquid suspension and on surfaces, there is a notable lack of standardized testing methods for antimicrobial products intended to be used against pathogens in the air (27). Moreover, even well-established surface testing methods may not be directly relevant for aerosol-transmitted pathogens, as they fail to capture the dynamics of airborne exposure and transmission risk. The exception to this for airborne testing is ASHRAE 185.3 (28) standard that defines the testing methods for evaluating commercial and industrial in-room air-cleaning devices and systems specifically for their effectiveness in removing or inactivating microorganism bioaerosols in a test chamber. Specifically, for airborne antimicrobials to be effective, they must directly contact pathogens in the air—either as vapor or aerosolized droplets. When a liquid antimicrobial product is tested for efficacy, it may be sprayed as large droplets or wiped onto a surface (coupon) or evaluated using suspension-based methods. In these cases, the product typically comes in contact with either the dried form of the microbe or interacts directly with microbes in a liquid suspension or wet droplet form. Unlike surface-based applications, where products interact with dried microbes, airborne delivery involves the dynamic interaction of the agent with suspended microorganisms. Our findings with TEG highlight that the mode of application significantly influences inactivation efficacy and time, reinforcing the need to evaluate antimicrobial products under conditions that reflect their intended use. In addition, aerosolizing test organisms allows for a more realistic assessment of antimicrobial efficacy by accounting for environmental factors like humidity and temperature that influence microbial susceptibility. Compared to surface-bound microbes, aerosolized organisms better represent real-world exposure to airborne pathogens. Moreover, dried surface samples can artificially shield microbes, potentially underestimating a product’s effectiveness in air.

The collected data across multiple trials in this study consistently demonstrate that TEG delivered by Grignard Pure™ exhibits significantly greater antimicrobial efficacy in aerosolized applications compared to hard surface treatments, underscoring the critical importance of evaluating air treatment products under conditions that closely mimic their intended real-world use. Across diverse microbial groups—including Gram-positive and Gram-negative bacteria, mycobacteria, viruses, and fungi—the aerosolized treatment of airborne microbes with TEG delivered by Grignard Pure^TM^ consistently achieved higher and more rapid reductions in viable organism concentration than surface treatment by the same product in liquid phase (Figures 2 and 3), where reductions were slower and less pronounced. Treatment of microorganisms on surfaces by TEG vapor delivered by Grignard Pure™ was even less effective (Figure 4). These findings are corroborated by testing conducted by the EPA’s Office of Research and Development (EPA ORD), which similarly reported superior log₁₀ reductions in airborne pathogen concentration following aerosolized TEG application compared to surface treatment, further validating the notion that the efficacy of antimicrobial products intended for air treatment should be tested in airborne phase (14) and not on surface-bound microorganisms. By the same token, the data suggest that TEG will not likely eliminate the need for surface sterilization, which is important to minimize the transfer of fomite-mediated infections. However, the use of a combination of TEG for air treatment and specific agents, e.g., isopropyl alcohol, for surface treatment is worth exploring in a separate study for potential synergistic effects.

On the other hand, the antimicrobial products designed for surface inactivation might not be effective in the airborne phase. A study conducted in 1963 examined the comparative inactivation of a germicide-resistant strain of phage *S. cremoris* 144F by surface-active compounds applied as aerosols. The study observed that surface-active compounds such as quaternary ammonium compounds (QAC) (tested in airborne concentrations of 0.024 ppm and 0.097 ppm) and phosphoric acid wetting agent (PAWA) (tested in airborne concentrations of 0.024 ppm, 0.048 ppm and 0.097 ppm) were ineffective as aerosols, despite their virucidal activity in aqueous solutions (29). Conversely, in a separate study, PAWA, QAC, hypochlorites, and iodophors, which are commonly used as sanitizing agents in the dairy industry, were tested for their virucidal efficacy in aqueous solutions against the same *S. cremoris* 144F. The PAWA sanitizer at a concentration of 12.5 ppm, as well as QAC at a concentration of 50 ppm completely inactivated the phage within 15 seconds (30). The inability of these compounds to act as effective agents in aerosol form can be attributed to the different physical environments under which they must exert their activity (29).

Thus, the data from the research described in this paper, as well as existing literature, strongly support the need to test antimicrobial products in the environment they are designed to treat. Otherwise, extrapolating surface disinfection data to predict airborne efficacy or extrapolating airborne efficacy to predict efficacy on surfaces might lead to inaccurate, and even diametrically opposite conclusions.

Another aspect to consider in developing and adopting testing protocols for airborne disinfectants is the sequence of introducing the inactivating product vs. introducing test microorganisms. The EPA ORD study showed that the sequence of product introduction — whether MS2 bacteriophage was introduced into a TEG-saturated chamber or vice versa— strongly influenced early kill kinetics. Introducing MS2 into a TEG-saturated chamber resulted in up to an additional 1 log₁₀ reduction within the first 15 minutes, compared to adding TEG to a virus-laden chamber (27). However, data sets from both test scenarios demonstrated that aerosolized TEG treatment delivered by Grignard Pure™ at concentrations of 1.2–1.5 mg/m³ achieved 2–3 log₁₀ reductions in airborne MS2 within 30–90 minutes. In our view, if TEG is eventually approved for use as an aerosolized antimicrobial product, it should be prophylactically deployed, i.e., people who may be infectious will enter spaces that have an effective preexisting level of TEG in the air. Based on the earlier mentioned research of ORD (14), as well as our earlier study (5), prophylactic use would produce greater reductions in the levels of airborne pathogens than releasing TEG only after infected individuals begin to sneeze or cough. Thus, we submit efficacy trials should be designed to reflect the real-world use of TEG and should release the inactivating product into a test chamber before introducing the test microorganisms.

Thus, an understanding of how various factors might impact the observed product efficacy is necessary for predicting performance in the real world and is important for developing useful and reliable standardized test methods (27). Ultimately, the most accurate assessment of an antimicrobial product’s efficacy requires a test methodology that mirrors its real-world application. For air treatment products, this means introducing both the test mixture and the test organisms into the air during efficacy studies.

Another critical aspect of developing a proper testing protocol involves selecting appropriate test microorganisms, especially if a product will be used against BSL-2 microorganisms or higher. Testing against such microorganisms in the airborne phase represents substantial technical obstacles, e.g., the need for at least BSL-3 testing environment, which cannot be surmounted by smaller companies or commercial testing laboratories. Therefore, the use of valid BSL-1 surrogates in such testing is critically important so long as data could be extrapolated to more pathogenic BSL-2 or even higher counterparts. When properly selected, BSL-1 surrogates provide a scientifically valid and practical alternative for evaluating airborne antimicrobial efficacy. Surrogates such as *K. aerogenes* and *P. fluorescens* have demonstrated reduction patterns on hard surfaces that closely align with those of selected pathogenic BSL-2 counterparts from the same genus (i.e., *K. pneumoniae* and *P. aeruginosa*), supporting their suitability for controlled testing in the airborne phase. Similarly, *S. epidermidis* serves as an effective surrogate for *S. aureus*. These correlations suggest that surrogate-based testing offers a framework for predicting the efficacy of antimicrobial products in real-world airborne scenarios, while maintaining safety during laboratory studies conducted at BSL-2 levels.

Additionally, this study uses BSL-1 Gram-positive and Gram-negative organisms as surrogates for airborne pathogens, but these groups also reflect skin- and enteric-derived contaminants. While aerosols may not address direct contact transmission, their use could help limit re-colonization of vulnerable hosts with antibiotic-resistant organisms in clinical settings—a potential application worth further study.

Although surrogate-based testing involves extrapolating performance from surrogate organisms to regulatory-recommended pathogens (e.g., US EPA), it provides a more scientifically defensible basis for assessing airborne efficacy than surface-only data. At the same time, we recognize that more comparative testing between the suggested surrogates (and perhaps other candidates) and their pathogenic counterparts might be needed, especially when developing and validating novel testing protocols, or when selecting surrogate microorganisms for testing of therapeutic agents. By validating air treatment products through aerosolized trials with appropriate surrogates, this methodology predicts that efficacy claims are both scientifically sound and reflective of real-world conditions. This approach addresses a critical limitation in traditional testing protocols and provides a reliable foundation for evaluating the potential of antimicrobial products to reduce airborne microbial contamination effectively. Insisting on airborne testing with microorganisms that cannot be studied in a BSL-2 facility would have the consequence of discouraging, if not preventing, the development of aerosolized antimicrobial products that could be effective tools for addressing diseases caused by transmission of airborne pathogens.

### Broad-Spectrum Efficacy of TEG-based product (Grignard Pure^TM^)

Our testing protocol aimed to evaluate the antimicrobial agent’s broad-spectrum potential under realistic conditions, including the use of Gram-negative and Gram-positive bacteria, non-enveloped and enveloped bacteriophages, mycobacteria, and mold. We have shown that TEG can achieve 2 to 4 orders of magnitude reductions in viable microbial counts of airborne bacteria, viruses, mold, and mycobacteria at total TEG concentrations (aerosol + vapor) ranging from 0.7 to 2.0 mg/m^3^ which are well within established safety margins – as discussed below in the “Safety of TEG” section. We also show that this reduction can occur in as fast as 35 minutes, with some organisms, like both tested bacteriophages, inactivated by almost two logs in 15 minutes. Two-log (99%) inactivation in 15 minutes or three-log (99.9%) inactivation in 35 minutes is equivalent to 18.4 and 11.8 air changes per hour (ACH), respectively (ACH is calculated using a first order kinetics equation C(t) = C(0) exp(-ACH x t), where C(0) is the initial microbial concentration, C(t) is the microbial concentration at time t in hours). This is substantially faster than airborne microorganism removal by ventilation systems operating at 5 ACH, a current CDC recommendation (https://www.cdc.gov/niosh/ventilation/prevention/Aim-for-5.html). Sultan et al. recently further confirmed that aerosolized TEG delivers consistent antimicrobial activity against MS2 bacteriophage across a broad range of real-world HVAC conditions—including temperatures of 22–25 °C, relative humidities of 40–70 %, air recirculation rates from 0 to 6 ACH, outdoor ventilation from 0.8 to 5 ACH, and filtration efficiencies ranging from no filter to MERV 14—demonstrating that aerosolized TEG maintains high inactivation rates even under dynamic ventilation and filtration scenarios (15). Notably, Sultan et al.’s study used lower total (aerosol + vapor) TEG concentration of 0.44 mg/m^3^, which could explain why some of the inactivation values in their study were lower than those observed in this study, where somewhat higher TEG concentrations were used.

The data presented by our study also suggest that Gram positive bacteria, both tested bacteriophages, and mycobacteria are highly susceptible to airborne treatment by TEG, while Gram-negative bacteria such as *S. typhimurium* and *P. fluorescens* are susceptible but not to the same extent. Fungal spores of *A. brasiliensis* have proven to be the hardiest, but even they were inactivated with ∼95% efficiency after 35 minutes of exposure.

These results emphasize the importance of testing antimicrobial air treatment products against diverse microorganisms to ensure comprehensive assessments of their real-world performance in the airborne phase. Notably, among the organisms tested, viruses and mycobacteria—those most strongly associated with clinically significant airborne transmission—showed substantial susceptibility to TEG, highlighting its potential impact where it is most needed. The data also corroborate the efficacy and utility of a TEG-based reagent (Grignard Pure^TM^) to decontaminate an indoor air environment and should provide a high degree of protection against airborne transmission of a number of different pathogens.

### Mechanism of Action

To understand the reason for the higher efficacy of TEG in air treatment compared to surface treatment, we hypothesize that when certain compounds are atomized and vaporized, they not only vaporize but also create microdroplets with a high surface-to-volume ratio that supports the efficient evaporation of active molecules to maintain vapor saturation. The exact mechanism behind TEG’s higher efficacy in air treatment remains under study. Studies suggest that TEG vapor, due to its high diffusion coefficient, may readily contact airborne microbes. As Puck et al. demonstrated (3), TEG vapor can condense onto microdroplets carrying microorganisms, achieving lethal local concentrations more rapidly than liquid-phase treatments. These observations point toward a vapor- and condensation-driven mode of action. Its hygroscopic nature may further contribute to microbial inactivation upon condensation (8). In addition to membrane permeabilization, proposed in earlier literature, TEG may additionally act by altering water activity, protein hydration and structure, as suggested for inactivation of non-enveloped viruses (13). Further studies are warranted to confirm the multiple mechanisms of action of TEG against different classes of microbes.

### Public Health Significance

For an airborne antimicrobial to be effective, it must directly contact pathogens in their most relevant form—suspended in air as vapor or microdroplets. Unlike surface-based methods, which can artificially shield microbes and misrepresent product efficacy, aerosolized testing offers a better representation of the true dynamics of airborne transmission. Our findings demonstrate that TEG performs optimally under these conditions, inactivating a broad spectrum of pathogens within minutes, and at concentrations well below the ACGIH (American Conference of Governmental Industrial Hygienists) safety threshold of 10 mg/m³ (https://www.acgih.org/triethylene-glycol/).

Safety of triethylene glycol has been extensively studied and a comprehensive discussion on the safety of TEG is presented in Desai et al. (5). TEG exhibits very low toxicity through oral, dermal, and inhalation exposure routes, with no toxicological endpoints of concern identified (31). As described by Desai et al. (5), the U.S. EPA classifies TEG as a "Safer Chemical" and has exempted it from tolerance requirements for antimicrobial residues on food-contact surfaces. Long-term inhalation studies in animals have shown no systemic adverse effects at concentrations significantly higher than those used in Grignard Pure™. A Margin of Safety analysis conducted by Nelson Labs further concluded that continuous inhalation exposure to TEG from product use presents a low risk of adverse effects, with no expected concerns related to acute, subchronic, or chronic toxicity, genotoxicity, or carcinogenicity (32). Moreover, TEG has been safely utilized in lighting effects (https://entertainingsafety.com/knowledge-base/understanding_fog_effects/) and air fresheners (https://edu.rsc.org/news/air-fresheners/2020753.article) for decades without significant health issues.

These results carry important implications for public health. During the COVID-19 pandemic, public health responses largely relied on personal behaviors—masking, distancing, and vaccination—despite inconsistent scientific consensus and variable compliance. This approach, though necessary, proved insufficient in stopping airborne spread and contributed to erosion of public trust. According to the hierarchy of controls, as explained by the CDC (https://www.cdc.gov/niosh/hierarchy-of-controls/about/index.html), engineering controls are strategies that reduce hazards from coming into contact with individuals. Our data demonstrate that TEG functions in this category by rapidly inactivating non-pathogenic airborne surrogates, consistent with historical studies showing that TEG vapors inactivate pathogens such as *Staphylococcus aureus*(8) and influenza viruses (7), indicating its broader relevance for airborne infection control. Further, based on the inactivation of a broad range of BSL-1 surrogates in the air and results from testing surface-bound BSL-2 organisms, we posit that TEG as delivered by Grignard Pure™ will also be efficient against airborne BSL-2 organisms.

### Study limitations

Our findings demonstrate the substantial antimicrobial efficacy of triethylene glycol (TEG) against a diverse array of airborne and surface-bound microorganisms; however, several limitations of the study should be acknowledged. Although this work reports many replicates with respect to the efficacy of TEG per se, inactivation of all organisms, except for MS2, was only tested once. Nevertheless, the inactivation kinetics were consistent across organisms, reaching net inactivation of 2-3 logs for bacteria and viruses. The consistent results allowed us to calculate average values and uncertainty (e.g., standard deviations) for treatment of related organisms, as shown in Figs. 2 and 3. Moreover, in Figure S1 we compare the average inactivation of different biological entities - bacteria (n=6), viruses (n=2), and a fungus - demonstrating 2-3 log inactivation of bacteria and viruses at the sampling time marks of 35 min and later. The inactivation uncertainty for different biological entities is also within relatively tight bounds (except for the 10-minute treatment of bacteria). Thus, consistent inactivation results obtained with multiple organisms comprise a "de facto" aggregate replication, demonstrating the efficacy of TEG to inactivate a variety of airborne pathogens. The rapid rise in inactivation of bacteria and viruses within the first 10-15 minutes also invites future investigation of the inactivation kinetics on this time scale more closely. Higher initial concentrations of airborne microorganisms would also allow for more robust determination of time-dependent inactivation kinetics.

Additionally, all studies were conducted within a controlled range of temperature and relative humidity (RH). Both temperature and RH can affect aerosol dynamics, droplet evaporation rates, and microbial viability, which in turn can influence the germicidal activity of vapor-phase agents such as TEG. Previous research has shown that TEG vapor exhibits optimal antimicrobial efficacy at mid-range RH (approximately 50–60%) and moderate temperatures (∼22 °C), with reduced performance observed at higher or lower RH or elevated temperatures (15). Moreover, environmental fluctuations in temperature and RH — such as those encountered in real-world indoor settings—might alter aerosol persistence and microbial susceptibility (33, 34). Therefore, subsequent studies should systematically vary temperature and RH to better assess the germicidal efficacy of TEG against various microorganisms.

Lastly, another limitation of this study is the absence of a representative soil load in the aerosolization medium. In real-world settings, airborne pathogens are often embedded in organic and non-organic matter or biological fluids, which can reduce the effectiveness of antimicrobial agents (35). Future studies should incorporate soil loads, such as proteins or mucins, to better simulate real-world bioaerosols and evaluate TEG’s performance under more realistic conditions.

### Data availability

Data presented in this manuscript are available upon request.

## ACKNOWLEDGMENTS

The authors acknowledge with gratitude helpful discussions and advice from Toni K Choueiri, William Esposito, Andre Fay, Antony Galione, Altaf Lal, and Donald W Schaffner.

Some of the research described in this article was funded by Bleu Garde, LLC, the company that makes Grignard Pure™.

